# Conditioning on a collider may induce spurious associations: Do the results of Gale et al. (2017) support a protective effect of neuroticism in population sub-groups?

**DOI:** 10.1101/231969

**Authors:** Tom G. Richardson, George Davey Smith, Marcus R. Munafò

## Introduction

Gale and colleagues (Gale et al., 2017) examined the association between neuroticism and mortality in a large sample (N > 300,000) drawn from the UK Biobank study (Sudlow et al., 2015). They observed that neuroticism was associated with an increase in all-cause mortality, but that following adjustment for self-rated health neuroticism was associated with a reduction in all-cause mortality. Further analyses stratified on self-rated health suggested that higher neuroticism was associated with reduced mortality only among those with fair or poor self-rated health. The authors conclude that neuroticism may have protective effects among certain sub-groups, and finding that generated substantial interest (TIME, 2017).

The availability of very large cohort studies such as UK Biobank in principle allows to identify associations where the absolute effect size may be small, but population-level impact considerable (as is the case of the results reported by Gale and colleagues). However, when two variables independently influence a third variable, and that third variable is conditioned upon, this can induce collider bias, which can distort observed associations (Munafo et al., 2017).

In the case of neuroticism, self-reported health and mortality, it is plausible that both neuroticism and risk factors associated with risk of all-cause mortality might influence self-reported health. In that case, conditioning upon self-reported health might induce collider bias, and generate spurious or distorted associations between neuroticism and both risk factors associated with all-cause mortality and all-cause mortality itself.

We explored this possibility using the same sample drawn from UK Biobank as that used by Gale and colleagues. Specifically, we examined the association between neuroticism and a range of risk factors known to be associated with all-cause mortality, both unstratified and stratified by self-reported health.

## Methods

We reproduced the analyses reported by Gale and colleagues as closely as possible using data derived from the UK Biobank study. A full list of the variables we used can be found in Supplementary Table 1. To verify that our dataset was similar to the one analysed by Gale and colleagues, we used Cox proportional-hazards regression to reproduce the hazard ratios for all-cause mortality in all individuals and within each self-rated health strata as reported by their study.

Linear and logistic regression were used to assess the relationship between neuroticism score and each covariate in turn (as shown in Table 1 in the study by Gale and colleagues) for continuous and binary traits respectively, with adjustment for age and sex. Analyses for each covariate were then repeated after stratifying individuals according to their self-rated health status. Our interest was in the comparison between the unstratified and stratified analyses.

## Results

We were able to reproduce the observations reported by Gale and colleagues when evaluating the relationship between neuroticism and mortality (Supplementary Table 2). Specifically, we observed hazard ratios < 1 and p values < 0.01 in the ‘Fair’ and ‘Poor’ selfreported health strata. These hazard ratios were interpreted in study by Gale and colleagues as neuroticism having a protective effect on mortality for individuals who classed themselves as having ‘Fair’ or ‘Poor’ health.

However, we also observed evidence suggesting that conditioning on self-reported health status may strongly influence the relationship between neuroticism and other risk factors in this study (Supplementary Table 3). In particular, we observed an instance of Simpson’s Paradox (Simpson, 1951) when assessing the relationship between neuroticism and body mass index after stratifying by self-reported health status. Specifically, we observed a negative association between neuroticism and body mass index in every stratum, but a *positive* association in the unstratified analysis. These results are shown graphically in Fig. 1. Examples of collider bias were also observed in the analyses with forced expiratory volume, cancer and diabetes (see Supplementary Table 3).

**Figure 1.**
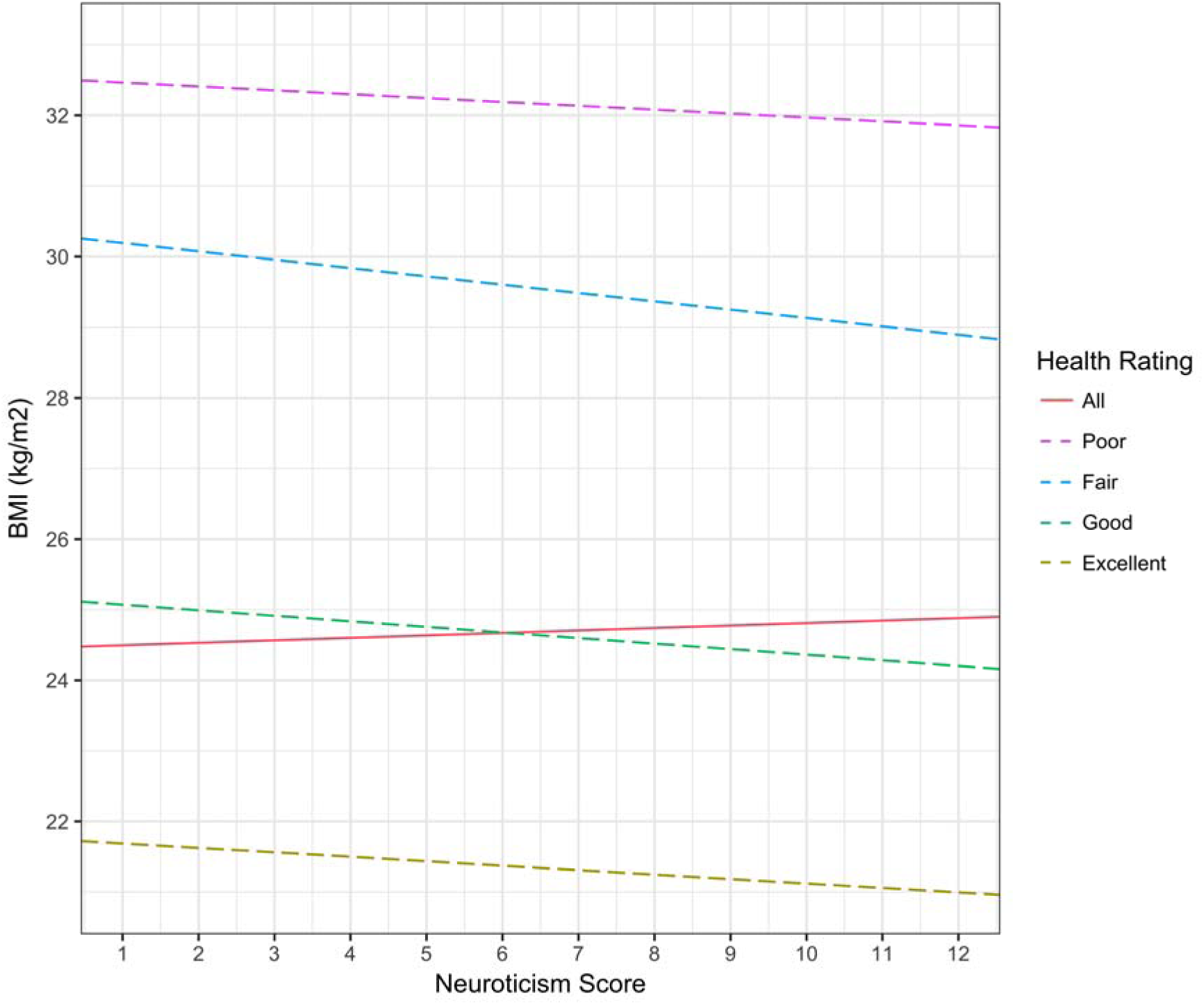
An illustration of potential collider bias in the UK Biobank study when assessing the relationship between neuroticism and body mass index.

Regression lines from the analysis between neuroticism and body mass index in the UK Biobank study. We observed a positive association in the unstratified analysis (i.e., all individuals, shown in red), but the opposite when we stratified on self-reported health (i.e., a negative association in all strata).

## Discussion

Our results suggest that the findings reported by Gale and colleagues should be interpreted in the context of the potential for collider bias. Specifically, conditioning on self-reported health status strongly influences the relationship between neuroticism and a range of risk factors known to be associated. Two factors lead us to believe that these associations are spurious. First, for many risk factors (e.g., BMI) the associations are clearly negative in every stratum, but positive in the unstratified analysis, indicating that a form of Simpson’s Paradox is operating. Second, we do not consider it likely that neuroticism could have the kind of protective effect suggested by Gale and colleagues across *all* of the risk factors we observed (particularly given evidence of Simpson’s Paradox).

The results we observed could be due to a confounding effect of self-reported health status on neuroticism and other risk factors, or due to collider bias if neuroticism causes self-reported health status. Differentiating between these possibilities would require stronger evidence that neuroticism causes self-reported health status, for example using Mendelian randomization (Davey Smith and Ebrahim, 2003, Davey Smith and Hemani, 2014). It is also worth noting that collider bias can still influence the analysis of genetic factors, which are protected from some biases in observational studies but not from this form of bias (Munafo et al., 2017).

Overall, our results serve as a cautionary note that while large cohort studies provide unparalleled power to elucidate associations between risk factors and disease outcomes, the ability to detect ever smaller effect sizes increases the risk that relatively weak biases may distort our findings. In other words, with great power comes great responsibility.

## Acknowledgements

This work was supported by the Medical Research Council and the University of Bristol (MC_UU_12013/1, MC_UU_12013/6). M.R.M. is a member of the UK Centre for Tobacco and Alcohol Studies, a UKCRC Public Health Research Centre of Excellence. Funding from British Heart Foundation, Cancer Research UK, Economic and Social Research Council, Medical Research Council and the National Institute for Health Research, under the auspices of the UK Clinical Research Collaboration, is gratefully acknowledged. UK Biobank data was analysed as part of project 15825.

